# Identifying *C. elegans* Lifespan Mutants by Screening for Early-Onset Protein Aggregation

**DOI:** 10.1101/2021.12.14.472506

**Authors:** Daniel F. Midkiff, Adriana San-Miguel

## Abstract

Genetic screens have been widely used to identify genetic pathways that control specific biological functions. In *C. elegans*, forward genetic screens rely on the isolation of reproductively active mutants that can self-propagate clonal populations. Since aged individuals are unable to generate clonal populations, screens that target post-reproductive phenotypes, such as longevity, are challenging. In this work, we developed an approach that combines microfluidic technologies and image processing to perform a high-throughput, automated screen for mutants with shortened lifespan using protein aggregation as a marker for aging. We take advantage of microfluidics for maintaining a reproductively-active adult mutagenized population and for performing serial high-throughput analysis and sorting of animals with increased protein aggregation, using fluorescently labeled PAB-1 as a readout. We identified five mutants with increased aggregation levels, of which two exhibited a reduced lifespan. We demonstrate that lifespan mutants can be identified by screening for accelerated protein aggregation through quantitative analysis of fluorescently-labeled aggregates in populations that do not require conditional sterilization or manual separation of parental and progeny populations. We further analyzed the morphology of protein aggregates and reveal that patterns of aggregation in naturally-aging animals differ from mutants with increased aggregation, suggesting aggregate growth is time-dependent. This screening approach can be customized to other non-developmental phenotypes that appear during adulthood, as well as to other aging markers to identify additional longevity-regulating genetic pathways.

## Introduction

The process by which organisms undergo structural and functional decline with age is regulated by a complex network of genetic and environmental factors(*1*). Obtaining a greater understanding of the aging process becomes increasingly relevant to human health by the year, as a greater fraction of the world population reaches old age(*2*). A key first step in understanding the biological mechanisms that regulate aging is to identify the genetic pathways at play. The nematode *C. elegans* has been crucial to our current knowledge of genes that modulate longevity(*1*), and several approaches to search for lifespan-regulating genes have revealed key genetic pathways(*3*–*9*). Approaches for genetic screening that rely on novel phenotypes or the development of new tools provide an opportunity to probe an unexplored genotype-phenotype landscape and identify yet uncharacterized genes that regulate the aging process.

Several technical advantages make *C. elegans* an optimal model system to perform large-scale genetic screens of aging regulators, such as its short lifespan, genetic tractability, fast life cycle, and ease of culture(*10*). *C. elegans* is amenable to both forward and reverse genetic screens, which rely on random mutagenesis or targeted gene silencing, respectively. Large-scale reverse genetic screens can be carried out by feeding *C. elegans* populations a library of dsRNA-expressing bacteria(*11*). RNAi screens for longevity genes have revealed important pathways, such as those associated with metabolism and mitochondrial function, but require handling and monitoring large numbers of populations throughout their entire lifespan(*4, 5, 12*). These screens are typically carried out in animals sterilized through genetic changes or exposure to 5-fluorodexoyuridine (FUdR), to circumvent the need to separate adults from their progeny(*4, 5, 12*). While RNAi screens only allow for the assessment of the effects of gene silencing, forward genetic screens depend on random mutagenesis and can result in strong and diverse functional mutations. The first forward genetic screen for lifespan mutants relied on a temperature-sensitive mutant that is sterile at 25 °C (*7*), and thus allows monitoring population survival without progeny. Recently, a forward genetic screen for maximum population lifespan screened populations derived from individually mutagenized animals(*9*) using conditionally sterile animals. As with RNAi, these approaches require measurement of lifespan for clonal populations of each mutant to be analyzed(*7*), and are thus highly labor-intensive.

Traditional forward genetic screens rely on isolation of individuals from a mutagenized population, after which clonal populations can be generated and analyzed to validate the phenotype(*13, 14*). Since it is necessary to isolate putative mutants that are still reproductively active, this approach is not suitable to screen for lifespan-altering mutations. Alternative approaches have used correlates or predictors of aging as screening phenotypes. One study focused on the identification of animals with normal locomotion rates at the beginning of adulthood but reduced locomotion in the very late reproductive stages(*15*). Recently, a screen of animals in the late reproductive stage was performed by searching for individuals with increased levels of intestinal autofluorescence from lipofuscin(*16*), which is considered a marker of aging(*17*). The mutagenized population of animals was kept synchronized until the late reproductive stages by blocking embryonic development using FUdR. When worms were no longer exposed to FuDR, they resumed reproduction and allowed for progeny collection. Reproduction and FUdR exposure are known to drive changes in longevity under some conditions(*18*–*20*), and thus screening animals that can naturally reproduce could lead to the identification of thus far elusive aging genetic pathways. However, separating mutagenized parents from their progeny makes genetic screens at late reproductive stage prohibitively labor-intensive, and has thus not been possible before.

One characteristic displayed by aging individuals is increased levels of protein aggregation due to a decline in proteostasis(*21, 22*). Screens have used increased aggregation levels in aggregating PolyQ disease models to search for genes that regulate proteostasis(*23, 24*). However, protein aggregation has not been used so far as a marker for aging. One protein that has shown increased aggregation with age is the Poly(A)-binding protein PAB-1(*25, 26*). Unlike PolyQ, PAB-1 is expressed and aggregates naturally in wild-type *C. elegans*. Additionally, aggregation has been observed to greatly increase immediately following the reproductive period(*26*). We thus hypothesized that by screening randomly mutagenized animals for increased PAB-1 aggregation in the late reproductive stages, mutants with an accelerated rate of aging could be identified.

In this work, we developed a new approach to isolate *C. elegans* longevity mutants through forward genetic screens using protein aggregation as a marker for aging in naturally reproducing populations. We screened mutagenized animals at the late reproductive stage by integrating microfluidic systems for progeny removal and animal sorting, with quantitative image analysis and automated scoring of protein aggregation levels. From the individual animals sorted as positive for high levels of aggregation, we obtained a set of mutant strains with elevated aggregation in the late reproductive stages. We assayed the lifespan of each aggregation mutant and identified mutants with a significant reduction in lifespan. We have generated new mutant strains that exhibit reduced lifespan and increases in PAB-1 protein aggregation. We also demonstrate a new approach towards forward genetic screening for aging genes, while avoiding inhibition of reproduction, that is applicable to other aging correlates or late onset phenotypes.

## Results

### Quantitative Image Analysis of PAB-1 Protein Aggregation as a Marker for Aging

To reliably analyze the association between of PAB-1 aggregation and aging, and use it as a readout for genetic screening, we developed image processing algorithms that measure PAB-1 protein aggregates from fluorescence microscopy images in *C. elegans*. We used a previously developed transgenic line(*26*), where tagRFP::PAB-1 fusion protein is expressed throughout the pharynx. Previous work has shown that differences in aggregation levels between young and old animals are larger in the anterior section of the pharynx, which includes the first pharyngeal bulb(*26*), and we thus selected this region for quantification of aggregates. Each pharynx image was split into an anterior and posterior section by an intensity-based segmentation followed by identification of the two largest objects, which represented the anterior and posterior bulb (**Figure S1**). Identification of the anterior bulb was based on image orientation. Throughout this study, imaging was performed either on a straight channel of a microfluidic device, or on agarose pads on glass slides. For on-chip images, the worm orientation is consistent in the imaging channel, ensuring the anterior pharynx is the top object. In this case, the anterior and posterior regions are split by vertically bisecting the image at the midpoint between the two centroids. When images are acquired on a glass slide, varying orientations make distinction of the anterior and posterior bulb more challenging. In this case, we applied a similar horizontal or vertical image split at the midpoint of the centroids. The split image was then displayed to the user, who then provides input to specify which image contained the anterior pharynx. Aggregate quantification was thus restricted to objects identified within the cropped anterior pharynx image.

In young animals, PAB-1 is expressed broadly at a high intensity throughout the pharyngeal muscle tissue. Protein aggregates were distinguishable from diffuse localization by their relative higher intensity, sharp boundaries, and small size relative to that of the pharyngeal muscle. They are also distinct from the narrow fluorescence accumulation observed in the middle of pharynxes (potentially localized to pharyngeal epithelium, **Figure S2**). To quantify PAB-1 aggregates with these characteristics, we developed an algorithm (called *Screening Algorithm*) that binarizes images following a standard deviation filtering step (**Figure 1**). Since young, old, and mutant worms expressed PAB-1 at varying levels, it was necessary to account for differences in overall intensity between images. We controlled for these differences by scaling each image by the mean intensity of the pixels contained within the anterior object identified during the pharyngeal splitting step, prior to the standard deviation filter. Pharynxes also exhibited spatial differences in baseline intensity. To account for this variability, we then applied a stepwise segmentation process to identify objects of varying intensity, while discarding large, non-aggregate objects. At each step, the stringency of the segmentation threshold was increased and objects above a certain size were removed. The results of each step are combined in a binary image, which is then post-processed to remove long objects that appear in pharynx edges. As the threshold becomes more stringent, identified objects become smaller, and thus objects missed in early steps are captured in later thresholding steps.

**Figure 1.**
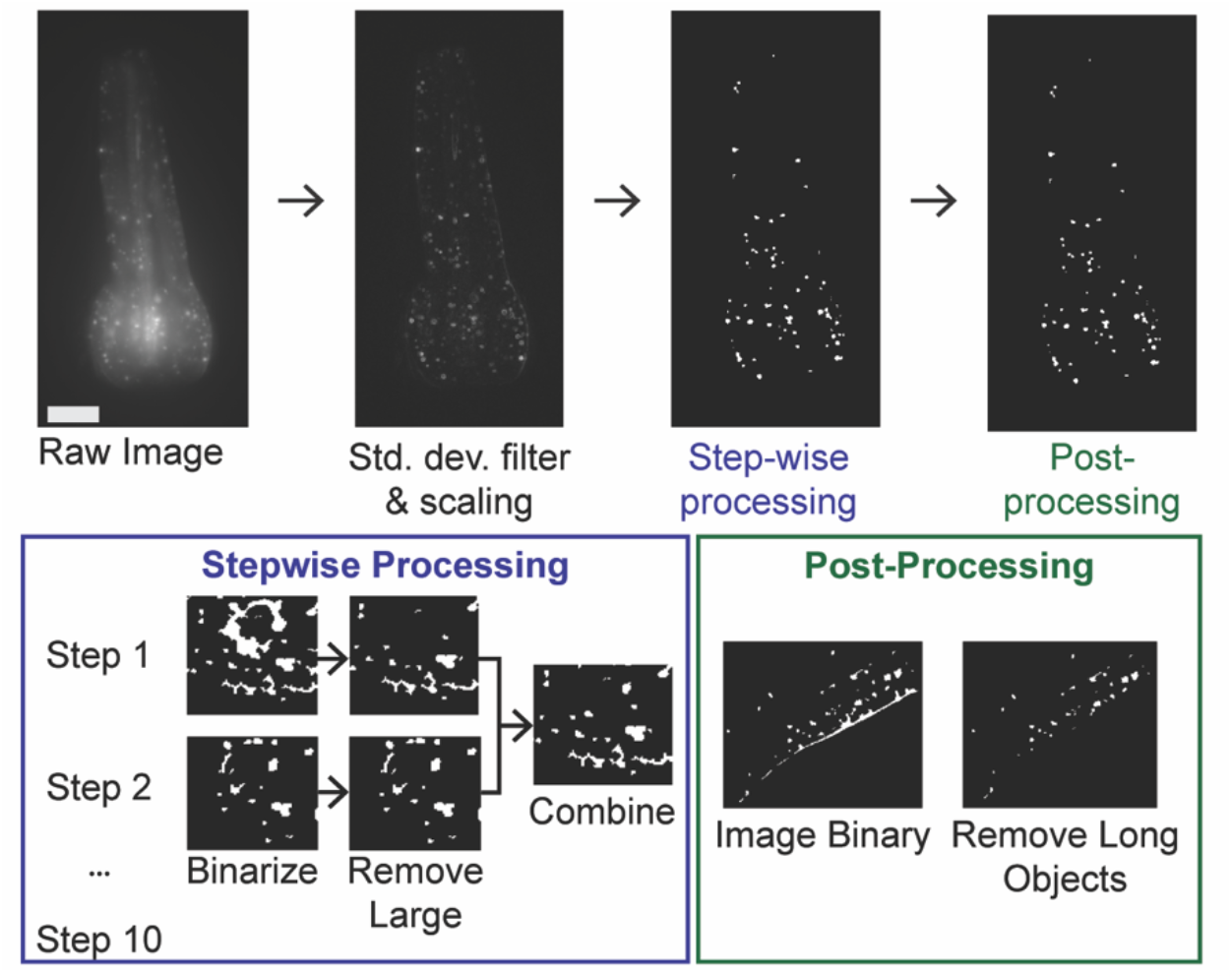
Quantitative analysis of PAB-1::tagRFP. Screening Algorithm image analysis flow: intensity-based scaling followed by step-wise aggregate identification and post-processing. Standard deviation of raw image acquired and scaled by the mean intensity of the anterior pharynx region. Image binaries acquired in a series of ten steps with an increasing threshold stringency. Step 1 is the least stringent threshold. In each step, objects above a defined size are removed. Objects that are missed in the initial step are added in subsequent steps, as they shrink below the aggregate threshold size. Images from each of the step are combined in an OR statement. Post-processing subtracts long objects to remove pharynx edges.

We validated the performance of this algorithm by comparing values calculated for total aggregate area and intensity to those identified through manual annotation. Our analysis of acquired PAB-1 aggregation images recapitulated previously observed trends in natural aggregation with age(*26*) and upon knock-down of the proteostasis regulator heat-shock factor 1 (*hsf-1)* (**Figure S3**). We thus determined that this algorithm could identify animals with high levels of aggregation, and thus used it for genetic screening, as described below. However, we found that the algorithm greatly underestimated the aggregation levels in images with high levels of diffuse protein, while performing better with images with low levels of diffuse protein (i.e. images from older animals) (**Figure S4A**). This trend was confirmed by plotting the total aggregate intensity determined by the algorithm versus the actual total intensity determined from the manual creation of ground truth binaries (**Figure S4B**).

To address these challenges, we next trained a supervised machine learning model (called *Validation algorithm*) to consistently identify aggregates with a large range of sizes and intensities. We created a training data set consisting of a sample of raw images and hand-drawn ground truth binaries. We defined the aggregate borders as the sharpest dividing line between the bright object interior and the baseline diffuse protein level. We extracted fourteen features from each raw image, using morphological operators designed to emphasize aggregates in images (**Supplementary Table 1**). The ground truth binary, the raw image (labeled Feature 1), and the thirteen other image features were used to train our classification models (**Figure 2A**).

**Figure 2.**
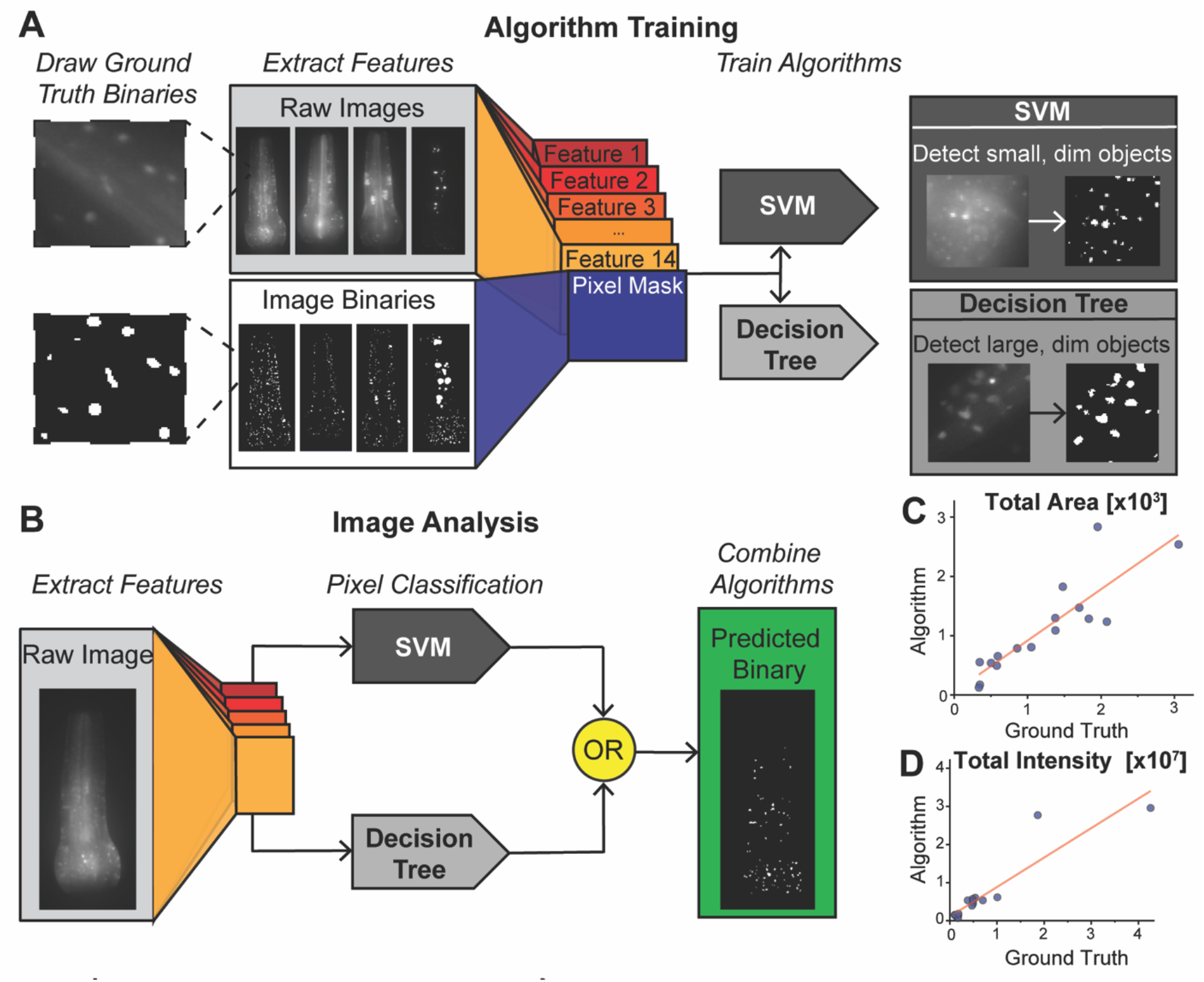
SVM and decision tree algorithms developed and trained to identify the presence of protein aggregates of PAB-1 in pharynx. Validation algorithm **(A)** Ground truth binary images. Features extracted from raw images to train pixel classification algorithms. Decision tree algorithm more accurately identifies large, dim objects, while SVM algorithm more accurately identifies small, dim objects. **(B)** Pixels in raw images are classified using both SVM and decision tree algorithm, two algorithms combined using OR statement. Validation ground truths constructed from 16 raw images. **(C)** Total aggregates area and **(D)** total aggregates intensity plotted for ground truth vs. algorithm classification(R^2^=0.76 and R^2^=0.84, respectively).

To ensure model robustness, we included images of pharynxes with different baseline intensities, and with aggregates of different sizes, shapes, and intensities. As would be expected, bright aggregates present in a dim intensity background are easily identified. However, aggregates that are only slightly brighter than the surrounding diffuse fluorescent intensity were detected at a lower frequency. Since each aggregate type exhibits different characteristics, we tailored two classification models to identify different aggregate types. By incorporating the results from both, this approach allows detection of a wide range of protein aggregate sizes and intensities. We trained a support vector machine (SVM) algorithm, which specialized in small and dim aggregates. We also trained a decision tree algorithm which was able to identify large aggregates present in dim images (**Figure 2A,B**). Bright aggregates were identifiable by both algorithms. The information from both algorithms was integrated by an OR statement, composing our classification algorithm.

To validate the performance of our classification algorithm, we constructed a validation set consisting of sixteen images and their corresponding ground truth labels generated by manual annotation. We quantified the performance of the algorithm by determining total aggregate area and total aggregate intensity. While some individual data points had a higher error, the algorithm-quantified aggregation levels in individual animals exhibited a strong correlation to the manually-annotated values (**Figure 2C,D**), which are also error-prone. To further assess the ability of this compound algorithm to measure protein aggregates, we imaged aging *C. elegans* populations at multiple days in wildtype and *hsf-1(-)* mutants, both previously shown to exhibit increased PAB-1 aggregation. The compound algorithm was able to recapitulate previously identified trends (**Figure 3**), further validating its applicability(*26*). These results indicate that the compound classification algorithm can reliably distinguish between the true levels of aggregate area and intensity throughout diverse images. As shown in **Figure 3**, this analysis reveals that age increases overall levels of PAB-1 aggregation, particularly at Day 7 once reproduction has stopped, when organismal fitness sharply decreases. In addition, significant levels of population heterogeneity are observed, pointing towards protein aggregation being subject to stochastic processes. As observed when comparing wildtype and *hsf-1(-)* populations, mutants exhibit much higher values of variability, indicating loss of *hsf-1* – regulated proteostasis is likely a driver of disordered protein aggregation. In addition, mutants also exhibit an increase in PAB-1 aggregation with age, suggesting *hsf-1* is not the sole regulator of aging-associated protein aggregation.

**Figure 3.**
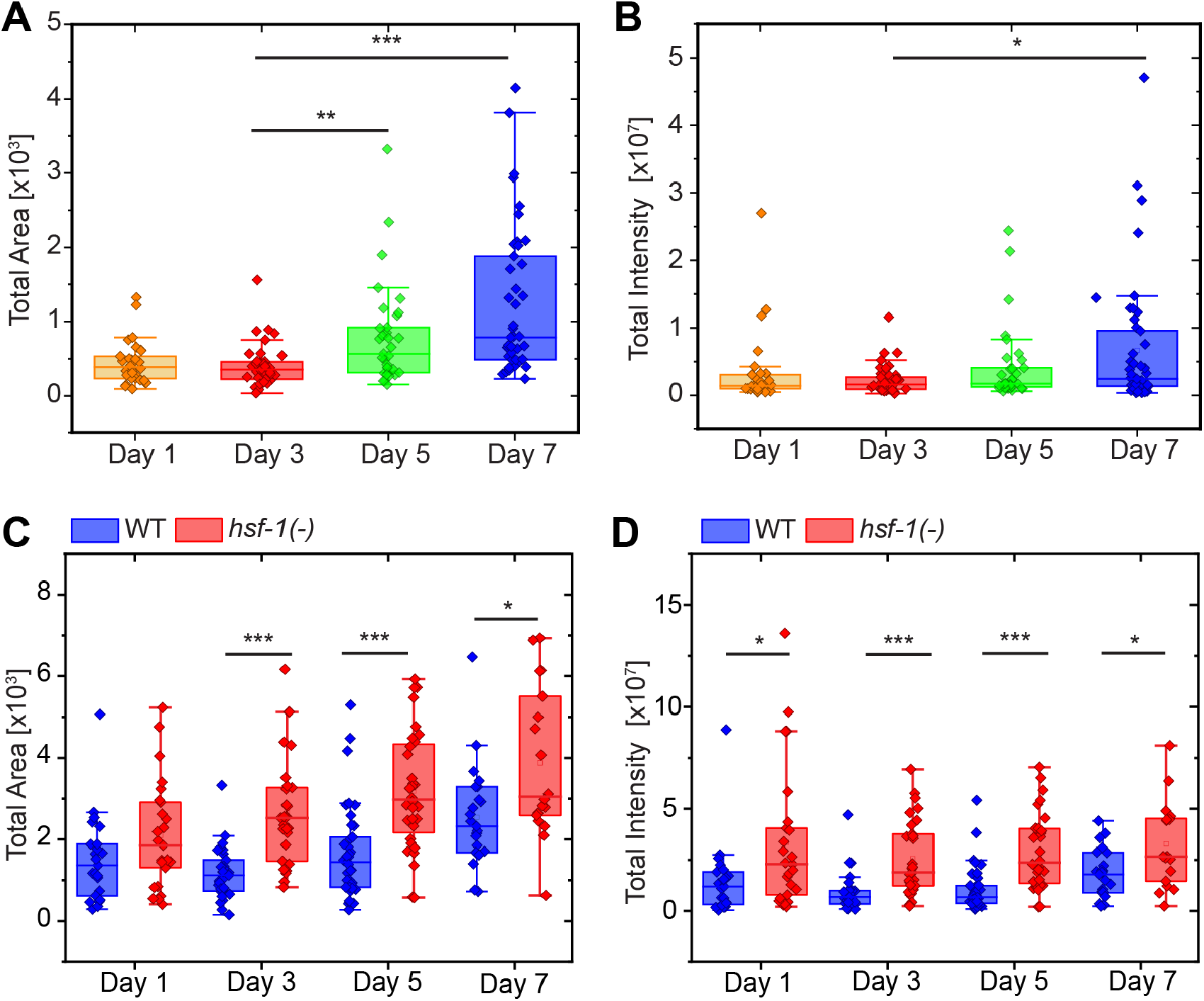
Increased levels of protein aggregation observed with age and in heat shock factor hsf-1 (-) mutants. Increase in total aggregate area **(A)** and intensity **(B)** observed from Day 3 to Day 7 of adulthood. Increase in total protein aggregation area **(C)** and intensity **(D)** observed in hsf-1 (sy411) mutant relative to wild-type transgenic strain *p<0.01, **p<0.001, ***p<1E-4 using Mann-Whitney U Test.

### Automated Genetic Screening for Increased Protein Aggregation in Reproductively Active Adults

Following the development of algorithms for identification of aggregation mutants, we performed a high-throughput forward genetic screen for increased PAB-1 protein aggregation. Previous studies indicate that PAB-1 experiences a spike in aggregation following the end of reproduction, and that mutations causing lifespan extension would produce a delayed onset of PAB-1 aggregation(*26*). We hypothesized that mutants that undergo aging at a faster rate would display increases in aggregation earlier in life. We observed an increase in aggregation beginning at Days 5 and 7 of adulthood in wildtype worms (see **Figure 3A,B**). With the goal of identifying animals with an early increase in age-induced protein aggregation, we chose to screen for increased PAB-1 protein aggregation at Day 3 of adulthood. Since animals start reproduction in Day 1 of adulthood, it is necessary to inhibit reproduction or to separate adults from progeny. In adult-based phenotypic analysis, inhibition of reproduction is typically performed through exposure to FUdR or starting with mutants that exhibit conditional sterility. We circumvent the need for inhibition of reproduction by the use of microfluidics approaches to separate randomly-mutagenized parent populations from their progeny. To achieve this, we utilized a microfluidic filtering chip we previously developed for long-term *C. elegans* imaging studies(*27*). The device consists of a microfluidic chamber surrounded by evacuation channels that allow embryos and larvae to exit the device while maintaining the synchronized adult population within (**Figure 4**). The device also has an attached array of channels for on-chip imaging, but this section was not used for the purposes of this study. A mixed population of parent worms and progeny (ranging from embryonic to 4^th^ larval stage) was loaded to the filtration chamber. Progeny was then removed by flowing buffer, leaving a population composed almost entirely of Day 3 adults. Following the filtration step, we used a microfluidic worm sorting platform previously developed for automated phenotypic screening in young adult worms (Day 1 of adulthood)(*28*). To screen larger Day 3 worms (70-80 µm vs. 50-60 µm in diameter), we developed a scaled-up design (**Figure 4, Figure S5**). We also adapted an imaging and screening software previously used for automated screening of synaptic patterning mutants(*28*). Prior to screening, the filtered adult population was exposed to 100 µM tetramisole for immobilization. Animals were then introduced into the worm sorting device, where individual animals are trapped, imaged, and sorted through pneumatically-operated on-chip valves. Control of the microfluidic system was performed with a programmable custom pneumatic box for fluid flow and valve operation. Putative mutants of interest are collected in one outlet for further analysis.

**Figure 4.**
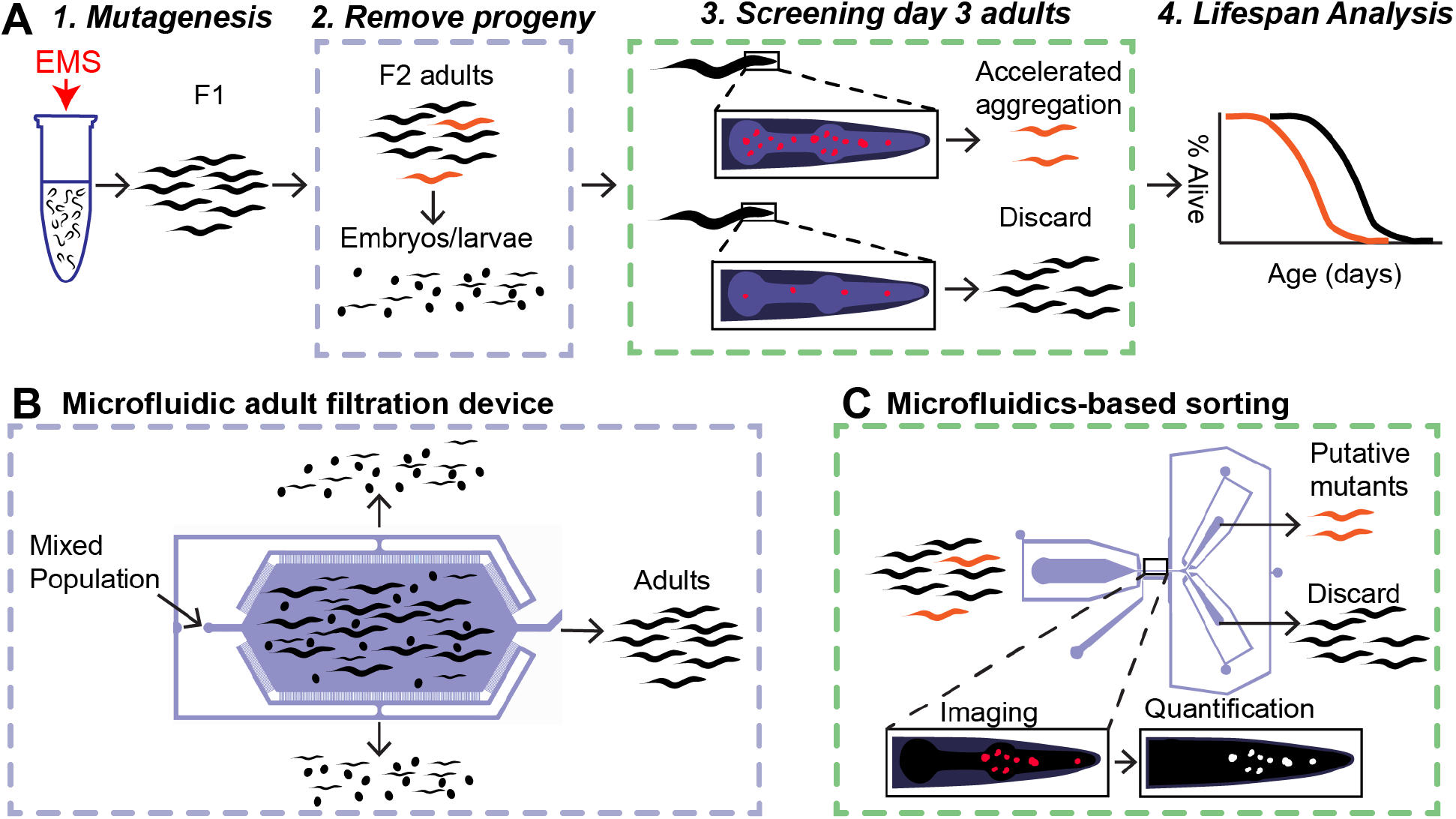
Process for semi-automatic forward genetic screening for protein aggregation mutants. **(A)** Initial parent (P0) generation of worms exposed to 100 mM EMS (Step 1). The population of second-generation progeny (F2) is grown to Day 3 of adulthood, where they undergo filtration to remove third-generation embryos and larvae (F3), producing a clean population of Day 3 adults (Step 2). Adult F2s undergo automated imaging and genetic screening in the microfluidic sorting chip (Step 3). Aggregation levels are quantified on-chip and animals with high levels of aggregation are isolated for follow-up analysis. **(B)** Microfluidic system for filtration of progeny of Day 3 adults. **(C)** Microfluidic worm-sorting chip for screening based on PAB-1 protein aggregation.

Animals were loaded, detected, imaged, and sorted in a high-throughput manner and without need for user interference. When worms were present in the field of view, they were identified through the baseline level of autofluorescence of a loaded animal, which triggers the closing of the inlet channel, as done in our prior work(*28*). Worms are randomly loaded into the imaging channel in one of two orientations: head-first or tail-first. Worm orientation was assessed through a mean intensity threshold, with head-first animals displaying a much higher mean intensity than tail-first animals, which were automatically discarded. Head-first worms were imaged by acquiring a z-stack, which was then compressed into a maximum projection for on-line analysis. Aggregation levels were then quantified on-chip through construction of an image binary using the Screening Algorithm. Animals were sorted as putative mutants if quantitative measurements exceeded a set threshold, which was set as the 99^th^ percentile of *C. elegans* aggregation levels in Day 3 wild-type adult populations. During the first portion of a screen, this value was set based on preliminary wildtype aggregation data of Day 3 adults. After 50 adults were screened, this value was reset based on a running calculation of aggregation data from the ongoing screen. Worms that were determined to exceed the aggregation threshold were sorted as positive and isolated for self-propagation. Of the 1902 animals that were screened, 60 were isolated as potential mutants. Of those, we recovered a total of 27 viable putative mutant strains that were subjected to further analysis. Some potential mutant worms were not recovered due to early ending of reproduction, mutations causing sterility, or being lost in transition out of the device.

### Identification of True Increased Aggregation and Lifespan Mutants

Once we isolated putative mutants from the genetic screen, we sought to verify the increased aggregation phenotype in clonal populations. Given the observed heterogeneity in aging populations, large data sets are required to confirm differences between wildtype and mutant populations. To minimize the number of large data sets that needed to be acquired, we performed this process in two steps. First, we acquired a small image data set from each potential mutant and wildtype, and quantified both total aggregate area and total aggregate intensity. We then calculated a two-sample z-score for each parameter and used this metric to rank strains (**Supplementary Table 2**). The putative mutant lines with the ten highest z-scores for each aggregation metric (area and intensity) were then selected for acquisition of a second, larger set of aggregation data (**Figure 5A,B, S6A**,**B**). From these analyses, we identified five strains that exhibited significantly higher aggregation levels than wildtype in either metric (**Figure 5A,B**).

**Figure 5.**
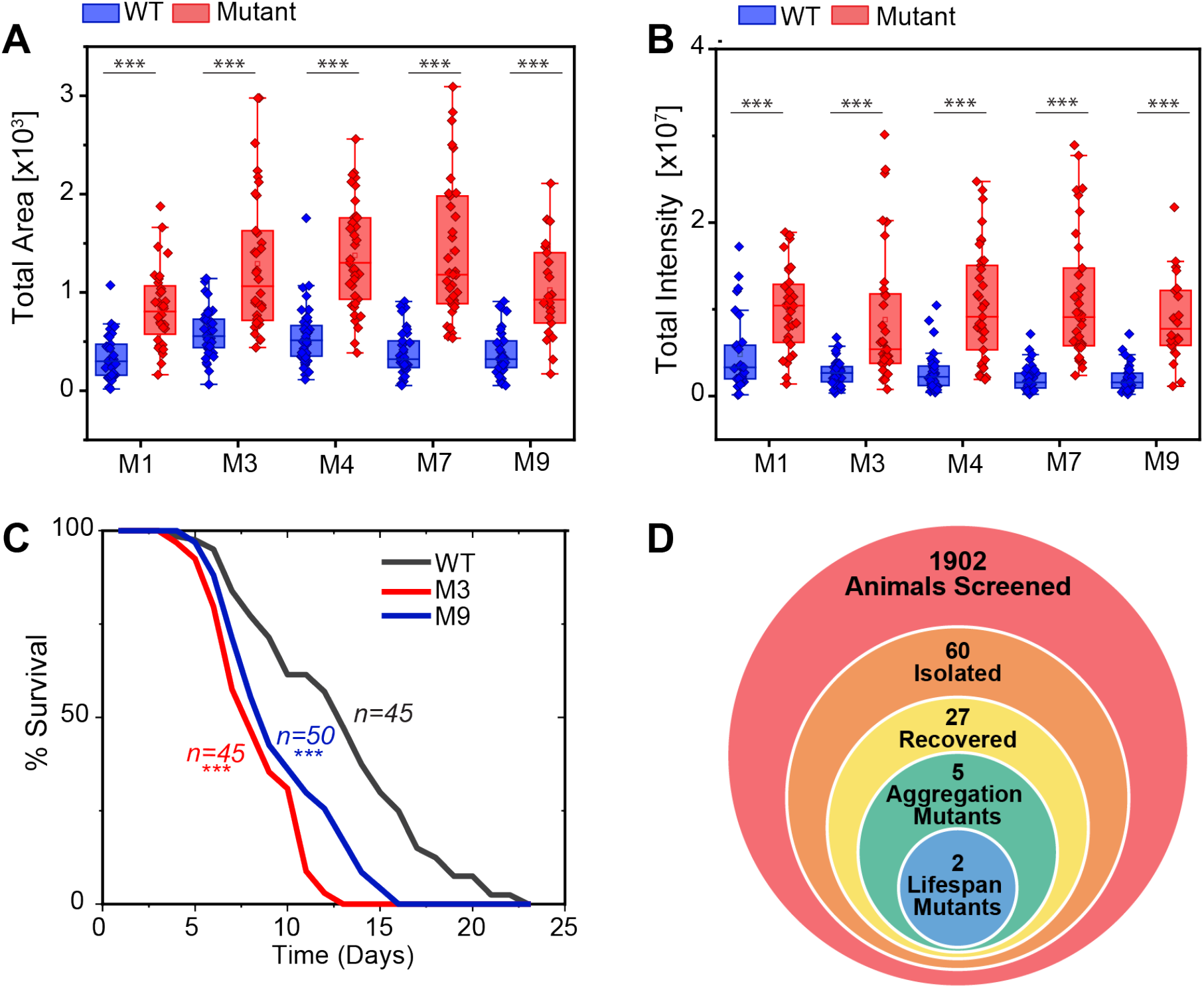
Verified mutants with increased protein aggregation as measured by: **(A)** total aggregate area, and **(B)** total aggregate intensity. **(C)** Mutants 3 and 9 also exhibit reduced lifespan. ***p<1E-4 using Mann-Whitney U Test for aggregation with Bonferroni correction, and log-rank test for lifespan. **(D)** Number of animals for each step in the screening process.

While finding mutants with increased levels of protein aggregation is of value, our overall goal is to identify mutants that carry a reduction in lifespan as well as an increase in aggregation. For each of the five verified aggregation mutants, we conducted a lifelong survival assay and identified two mutants with a reduced lifespan (**Figure 5C,5D**). Surprisingly, some mutants showed greatly increased aggregation but did not exhibit a corresponding decrease in lifespan (**Figure S6C**,**D**). In fact, the aggregation mutant with the highest z-score that we identified (Mutant 1), did not cause any lifespan alteration. It is thus not uncommon for mutants with strongly increased levels of PAB-1 aggregation to not carry a corresponding reduction in population lifespan. Previously published data has lent evidence towards a reduced level of protein aggregation being correlated with an extended lifespan(*26*). In the data we have acquired, we did not see this association in every mutant line, suggesting this association could only be present in long-lived mutants. However, it is also possible that increased aggregation in mutants that do not show disrupted lifespan could stem from mutations to tagRFP or PAB-1, rather than affecting proteostasis or longevity pathways. We also looked at known reduced lifespan mutants to determine if they exhibited an increased level of protein aggregation. Unlike *hsf-1*, which when knocked down was previously shown to reduce lifespan and increase aggregation(*26*), we observed no significant increase in aggregation in shortened lifespan *kat-1* mutants and a slight aggregation increase in *jkk-1* mutants(**Figure S7**). This aligns well with the prior observation that reduced insulin signaling reduces aggregate formation, as *jkk-1* positively regulates DAF-16 activity(*29*). These results suggests that protein aggregation increases with aging, but that genetic pathways that regulate aging do not necessarily modulate proteostasis.

We next aimed to assess whether the mechanisms driving aggregation in the identified mutants could be different, by analyzing aggregate morphology. Prior work found a significant increase in mean size with age, but this effect was not induced by rapid heat stress(*26*). To test if our identified lifespan mutants produced a similar increase in aggregate size, we examined the areas of a random sample of 100 aggregates from the wildtype and the five identified mutant strains at Day 3 of adulthood. We found that none of the mutants, regardless of any increase in lifespan, exhibited a significant increase in mean aggregate area (**Figure 6A**). In contrast, we did observe a significant increase in mean aggregate area for older worms (Day 7 of adulthood) as compared to late-reproductive worms (Day 3 of adulthood) (**Figure 6B**). We hypothesize that this increase in mean aggregate size is caused by the gradual growth of initially small aggregates through accumulation of aggregate protein over time. In contrast, in younger mutants, there is less time for aggregates to accumulate, though the propensity for PAB-1 to aggregate is high. Interestingly, we found that *hsf-1(-)* mutants on Day 3 of adulthood did produce a significant increase in mean aggregate size, albeit less drastic than for naturally aging worms (**Figure 6B**). This would indicate that some aggregation mutants can produce larger aggregates earlier in life. Potentially, this could indicate that the mutants identified from the screen do not cause a loss-of-function in *hsf-1*.

**Figure 6.**
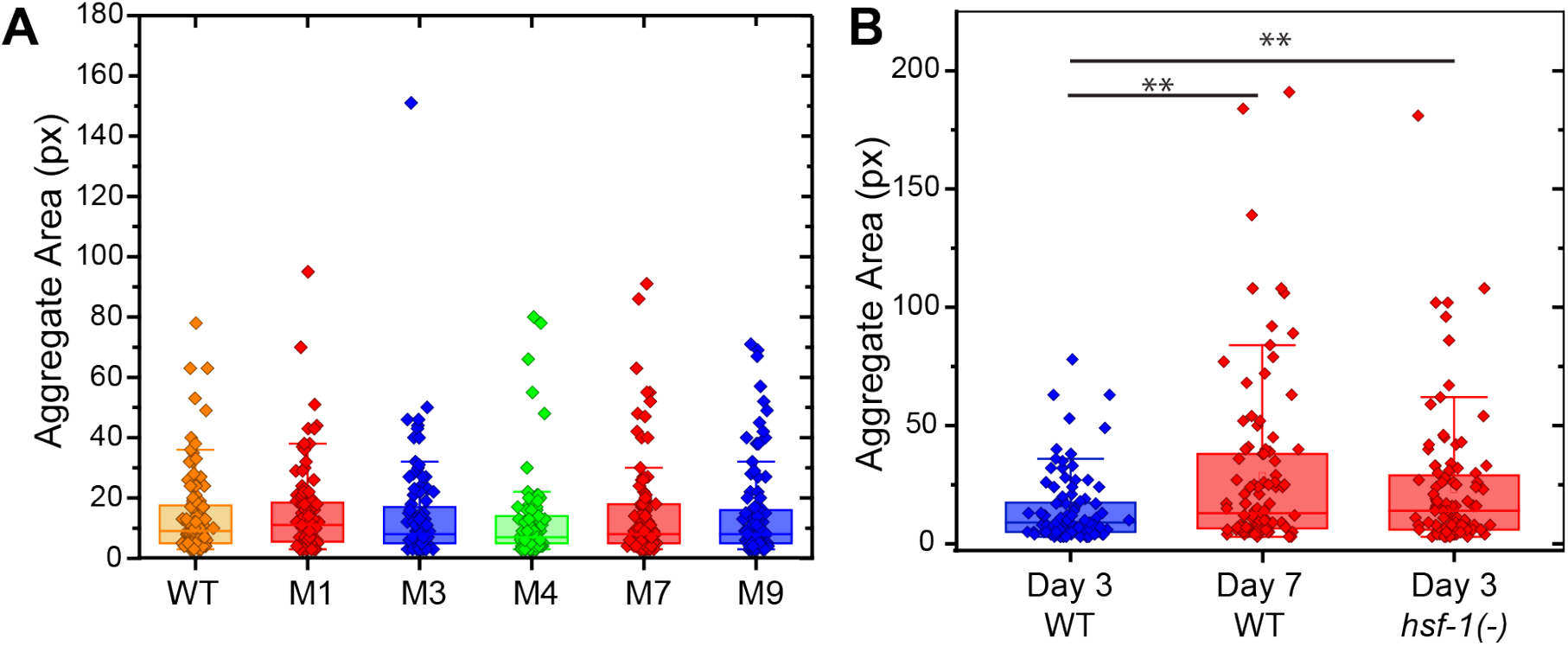
**(A)** Average area of a sample of 100 random PAB-1 aggregates from wild-type and each mutant with an increase in total aggregate area. No significant difference in size observed for any mutant. **(B)** Average area of a sample of 100 PAB-1 aggregates obtained from wild-type animals at Day 3, wildtype animals at Day 7, and hsf-1(-) mutants at Day 3 of adulthood. Aged and hsf-1(sy411) animals exhibit larger aggregates (** p<0.01 using Mann-Whitney U Test).

## Discussion

In this study, we employed image analysis and machine learning-based classification techniques to accurately quantify aggregation levels of the PAB-1::GFP reporter in diverse *C. elegans* populations. The largest challenge in accurately quantifying aggregation is defining the presence of an aggregate. We define protein aggregates visually as sharp puncta of fluorescence that are brighter than the broad level of protein expression in the pharynx. From these visual identification criteria, we were able to define image filtering operations that could identify the presence of protein aggregates based on the object size and intensity and the pharyngeal background intensity. While manual annotation is prone to subjective classification bias, the algorithms developed here can consistently identify aggregates in diverse image types. Being fast, accurate, and consistent, these methods enabled the identification of general increases in aggregation, and rapid, automated, on-chip quantification and sorting based on aggregation levels in mutagenized worm populations. Compounding the results of two machine learning techniques improved quantification of protein aggregation levels in dimmer regions of the pharynx, where aggregate identification is more challenging, and enabled identification of aggregates of different intensities and in images with a wide range of diffuse protein intensities and object sizes. To analyze the performance of our algorithms, we chose to calculate the areas and intensities determined from the classification algorithm (predicted) vs. the visually constructed ground truth (actual) for our validation data, since our goal was to use these metrics to differentiate between strains of baseline (wildtype) and increased aggregation. While we have trained our algorithm specifically for the PAB-1 protein, this method can be adapted to the quantification of other aggregating proteins.

In this study, we used PAB-1 aggregation as a proxy phenotype to detect lifespan mutants. Previous work reported trends of increased protein aggregation following the reproductive period, based on categorical classification of aggregation levels. Our analysis recapitulated this trend, and in addition enabled quantifying differences in protein aggregation beyond what was identifiable in categorical data sets. Using microfluidic technology, screening for increases in protein aggregation in a systematic manner without need for human decision making or operational interference was made possible. Moreover, in this work we developed a new method for identifying aging phenotypes in the late reproductive stage while circumventing the need for genetic or chemical sterilization, which could be a compounding factor on aggregation or the aging process(*30*). This method of screening opens up the possibility of searching for mutants that exhibit phenotypes late in life, without the need of temporary sterilization. Specifically for identification of lifespan mutants, this work shows that aggregation alone does not always correlate with longevity. Although we observed a sufficiently high rate of mutant identification, there are improvements that could be made to increase the rate at which lifespan mutants are identified, such as the simultaneous use of multiple markers of aging. We have shown that screening for aging phenotypes during the late reproductive period can effectively identify mutants with an altered lifespan. With the use of new or existing software for phenotype quantification, our method can be applied to other phenotypes related to aging. By conducting similar screens, the genetic networks that connect other phenotypes with the overall decline with age can be identified.

Screening of approximately two thousand animals yielded two lifespan mutants, a rate of 0.1%, while resulting in an aggregation mutant rate of 0.25 %. While we focused on the top ten highest aggregating mutants out of the 27 potential mutants recovered from the screen, it is possible that more aggregation or lifespan mutants could be identified upon further study of the remaining putative mutants. Amongst those studied, several strains cause an alteration in aggregation levels but not in lifespan. PAB-1 protein forms stress granules in response to heat shock(*26*), so it is possible that some of the identified mutations cause an alteration in heat shock response as opposed to an alteration in protein insolubility. However, heat and other stress responses have been associated with lifespan, though the correlation is not perfect(*31, 32*). Stressors can have opposing effects on lifespan: low levels can extend lifespan through hormesis(*33*), while high levels can be detrimental or even lethal(*32*). Likely, the association between aggregation and lifespan is dependent on environmental factors and exposure to stress, which could be explored in future work.

In this study, we have developed a method for identifying mutants with a reduction in lifespan by screening for mutants that undergo an increase in protein aggregation during the late reproductive stage. We developed a semi-automated pipeline that makes use of microfluidic lab-on-a-chip technologies to maintain age-synchronized parental worm populations which undergo automated aggregation quantification and on-chip sorting. This approach enables an otherwise unfeasible search that would require manual sorting of animals based on visual phenotypic classification. These experiments are highly intensive and require hours of manual labor to conduct. For PAB-1 protein aggregation and other high-resolution phenotypes, phenotypic quantification can be even more time consuming. Through further investigation of identified aggregation and aging mutants, we aim to identify genes and pathways which regulate *C. elegans* aging and proteostasis networks. Through increased understanding of the genetics of aging obtained through screening, there is potential to better understand how natural aging affects human health, and how humans can live longer and higher quality lives through advances in science and medicine.

## Limitations of the Study

In this work, we have developed software for assessing the level of aggregation within individual objects. As described in the Results section, the *Validation Algorithm* can robustly identify differences in aggregation between populations, and in many instances can distinguish between small total aggregation differences within individual animals. However, our algorithm is more limited in its ability to determine the true size of aggregates. We trained the classification algorithms using manually-annotated images, and although we pre-defined the characteristics of an aggregate, this could result in biased and inconsistent labeling. For instance, it was extremely difficult to consistently identify the exact location of the aggregate borders. In summary, this algorithm can be used to measure relative aggregation levels, rather than to quantify the exact amounts of aggregated protein. In addition, we performed our genetic screen with the less accurate *Screening Algorithm*, which could raise concerns of missed mutants with elevated aggregation. These concerns are rebutted by the high positive hit rate of recovered true aggregation mutants: out of 27 recovered mutant lines, five were determined to be true aggregation mutants. Smooth operation of the device required both functional optimization and practice. It was also necessary to account for disruptions in the operation of the device, most commonly due to clogging from small pieces of debris in the inlet media. For these instances, we incorporated an automated high-pressure flush cycle to dislodge the obstructing object. If these cycles were unsuccessful at dislodging the debris, a manual dislodge was necessary. Finally, as stated before, it is possible some of the identified mutations directly affected the propensity of PAB-1 or tagRFP to aggregate, rather than modulating organismal proteostasis or longevity pathways. Future genotypic analysis will enable addressing this question.

## Materials and Methods

### C. elegans strains

*C. elegans* strains DCD214 [*myo-2p::tagRFP::pab-1*], PS3551 [*hsf-1(sy411)*], KU2 [*jkk1(km2)*], and VS24 [*kat-1(tm1037)*] were acquired from the CGC (Caenorhabditis Genetics Center). Strains carrying integrated transgene *myo-2p::tagRFP::pab-1* as well as *hsf-1 (sy411), jkk-1 (km2)*, or *kat-1(tm1037)* were constructed by crossing DCD214 males with hermaphrodites from strains PS3551, KU2, or VS24 according to standard protocols. All other mutants were derived from ethyl methane sulfate (EMS) mutagenesis of the DCD214 strain.

### Worm Culture and Growth

Worms were cultured on NGM plates according to standard protocols(*34*) on lawns of *E. coli* strain OP50. Worms were grown at 20 °C, except for lifespan experiments, where they were grown at 20 °C until Day 1 of adulthood, and then transferred to 25 °C. Day 1 is defined as the first 24 hours following the conclusion of the L4 larval stage.

### Image Acquisition

Three different imaging settings were used for image acquisition. Setting A was used for the acquisition of all screening images, and the first round of aggregation data (**Supplementary Table 1**). Setting B was used for acquisition of the *hsf-1(-)* data sets (**Figure 2C,D**). Setting C was used to acquire the wild-type aging, *jkk-1*(-), and *kat-1(-)* data set, and the second round of aggregation data (**Figure 2A,B, Figure 4, Figure S7**). Image sets were scaled to the size and intensity of *Setting B* to account for differences between the settings as follows: Images acquired using Setting A were resized by a factor of 0.635 using the MATLAB “imresize” function to account for the change in magnification. The pixel intensity of images acquired using Setting C was multiplied by a factor of 0.5 to account for differences in mean image intensity. Images were acquired with each setting as follows:

#### Setting A

Images were acquired using a Leica DMi8 microscope and an Orca Flash 4.0 v3 CMOS camera. Fluorescent TagRFP protein was visualized at 630 nm wavelength and excited at a wavelength of 560 nm from a Lumencor Spectra X Light Engine, and filtered using a Leica TXR filter cube. Images were acquired at a magnification of 63x. During on-chip screening, images were acquired through immobilization of worms exposed to tetramisole in microfluidic channels. For initial aggregation assays, images were acquired by placing worms on pads made from 2% agarose dissolved in water between two glass slides, and were immobilized using 2 mM tetramisole dissolved in M9 buffer. Image z-stacks of 70 1-µm steps were acquired at an exposure time of 60 ms and were saved as a maximum intensity projection in a .tiff format. Image acquisition was controlled via a MATLAB custom graphical user interface.

#### Setting B

Images were acquired using a Leica DMi8 microscope and an Orca Fusion C14440 CMOS camera. Fluorescent TagRFP protein was visualized at 630 nm wavelength and excited at a wavelength of 560 nm from a Lumencor Spectra X Light Engine, and filtered using a Leica TXR filter cube. Images were acquired at a magnification of 40x. Images were acquired by placing worms on pads made from 2% agarose dissolved in water between two glass slides and were immobilized using 2 mM tetramisole dissolved in M9 buffer. Image z-stacks of 100 1-µm steps were acquired at an exposure time of 60 ms and were saved as a maximum intensity projection in a .tiff format. Image acquisition was controlled via a MATLAB custom graphical user interface.

#### Setting C

Images were acquired using a Leica DMi8 microscope and an Orca D2 CCD camera. Fluorescent TagRFP protein was visualized at 630 nm wavelength induced through excitation of samples from light at a wavelength of 560 nm from a Leica EL6000 metal halide external light source and filtered using a TXR filter cube. Images were acquired at a magnification of 40x. Images were acquired by placing worms on pads made from 2% agarose dissolved in water between two glass slides and were immobilized using 2 mM tetramisole dissolved in M9 buffer. Image z-stacks of 70 1-µm steps were acquired at an exposure time of 60 ms and were saved as a maximum intensity projection in a .tiff format. Image acquisition was controlled via a MATLAB custom graphical user interface.

### Image Processing

TIFF images were imported into MATLAB 16-bit integer arrays. Images that displayed blurred edges or asymmetric fluorescent protein localization were excluded from analysis. Images were split into anterior and posterior portions, and all image processing was performed on the anterior images (**Figure S1**). All images were processed through code developed in MATLAB. Image binary ground truths were created by manually drawing over aggregate objects using Fiji ImageJ analysis software. P-values for assays of total aggregation and mean aggregate size were determined using the Mann-Whitney U statistical test.

### Microfluidic Device Fabrication and Operation

Silicone master molds were fabricated in a cleanroom using SU-8 photoresist(*27, 28, 35*) spin-coated on silicon wafers. Master molds with multiple-layered features were fabricated through UV photolithography using an MA6 contact aligner, and unexposed photoresist was dissolved using 1-methoxy-2-propanol acetate. SU-8 mold features were exposed to vaporized silane following the development step. Device 1 refers to the device used for progeny filtration, and Device 2 refers to the device used for worm imaging and sorting. Microfluidic devices were fabricated using a mixture of SYLGARD 184 polydimethylsiloxane (PDMS) elastomer and crosslinker. A thin layer at the base of Device 2 was fabricated at a ratio of 20:1 elastomer base to crosslinker, while the remainder of Device 2 and all of Device 1 were fabricated using a 10:1 ratio. Mixtures of elastomer base and crosslinker were degassed in a vacuum-sealed chamber prior to molding to remove gas bubbles. Devices were cured by baking at 70 °C overnight, and then plasma bonded to glass slides using a plasma cleaner. Devices were operated using a controllable pneumatic pressure box built in-house and operated using MATLAB GUI software.

### Genetic Screening

Random mutagenesis was performed on a population of wild-type worms by exposure to ethyl methanesulfonate (EMS) using standard protocols(*36*). Worms were grown to the F2 generation following mutagenesis at temperatures ranging from 15 °C to 25 °C to stagger population growth. Worms were then age-synchronized using bleach solution (via standard protocols) and grown at 20 °C until the population reaches Day 3 of adulthood. Worms are washed from the plates using M9 buffer and loaded into the culture chamber of Device 1 (**Figure 4**). Flow was induced to flush progeny out of the device, leaving a population of exclusively Day 3 adults for screening. Adult worms were then partially immobilized using tetramisole (∼100 µM) and loaded into the inlet of Device 2 for on-chip imaging (**Figure S5**). Valve channels were filled with a 1:1 mixture of water and glycerol and operated using the pneumatic pressure box. A MATLAB graphical user interface was configured to control the device and perform automated screening. All code is available at Github (asanmiguel/AggregationScreening). As worms were loaded into the imaging channel of Device 2, the real-time detection of fluorescent light triggered the closing of the inlet valve. Z-stacks of immobilized were then acquired and compressed to a maximum projection image. Aggregation level was then quantified on-chip using code developed in MATLAB. Worms were sorted as potential mutants based on exceeding either a user-set total intensity value or total large intensity value (total intensity of objects that exceed a user-specified area). The threshold was set either by direct user input based on acquired experimental data, or by using the calculated 99^th^ percentile of data previously acquired in the ongoing screen. Sorted worms were diverted to one of two outlets by on-chip valves based on the measured aggregation level. Worms that exceeded the defined threshold were sent to a collection tube from where they were placed on individual growth plates where they produce putative mutant populations. Worms not sorted as putative mutants were discarded.

### Lifespan Analysis

Worms were age-synchronized and grown at 20 °C until reaching Day 1 of adulthood. Approximately 100 to 200 worms from each population were then counted and picked to a new plate and placed at 25 C. For each subsequent day, the number of living, dead, and censored worms were counted. Living worms are defined as worms that responded to a gentle touch from a platinum wire and dead worms are defined as those that did not respond to the same stimulus. Worms were censored if they were not visible (i.e. crawled off the plate or under the agar) or displayed signs of internal hatching. Signs of internal hatching include internal growth of larvae (“bag of worms”) as well as worms that undergo apparent rupture. Worms were picked to new plates daily during the egg-laying phase. Following egg-laying, worms were picked to new plates when plates begin to dry or in rare cases where small fungal contaminants appeared. After no living worms remained, mutants’ lifespan data was compared to wild-type data using the log-rank statistical test to determine P-values, and to identify worms with reduced lifespan(*37*).

## Acknowledgements

*C. elegans* strains were provided by the CGC, which is funded by NIH Office of Research Infrastructure Programs (P40 OD010440). This work was supported in part by the U.S. National Institutes of Health (NIH) grants R00AG046911 and R21AG059099.

## Author contributions

D.F.M. performed the experiments, developed image analysis algorithms, microfluidic devices, and analyzed the data. D.F.M. and A.S.M. conceptualized the work and wrote the manuscript. A.S.M. supervised the study.

