## Supplementary Figures and Tables for "Identifying *C. elegans* Lifespan Mutants by Screening for Early-Onset Protein Aggregation"

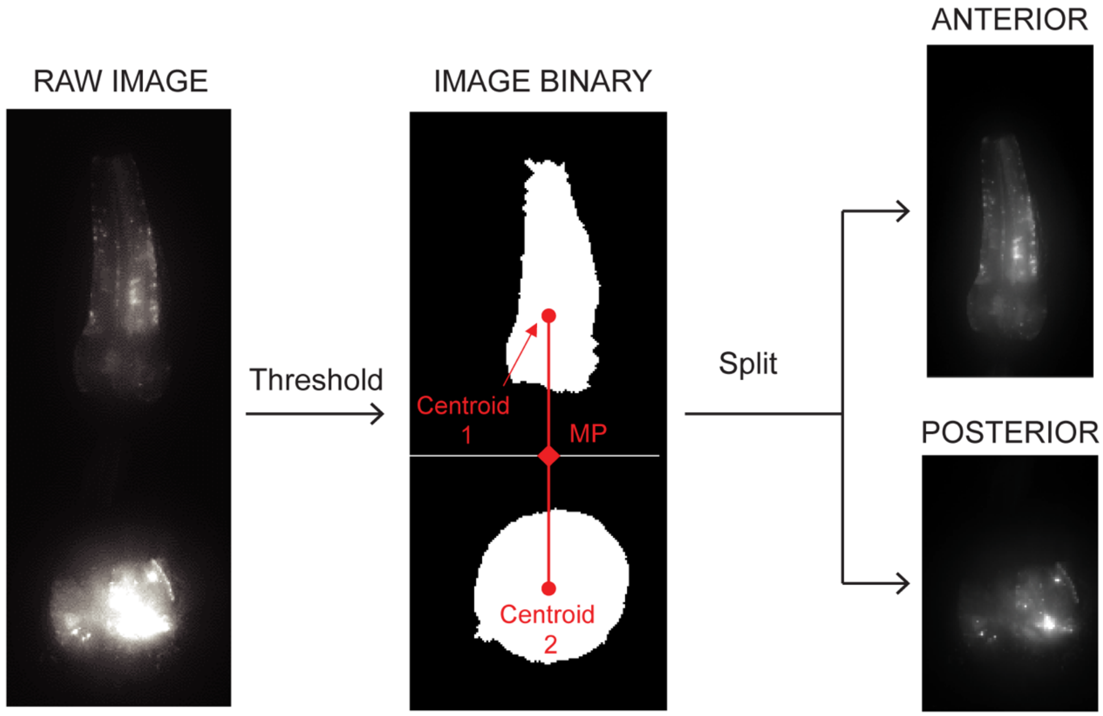


***Supplementary Figure 1*** *Pharyngeal image is split into anterior and posterior images. Binarization threshold applied to identify two largest objects, which are split at the midpoint of two centroids. Top of image is anterior, bottom is posterior.*

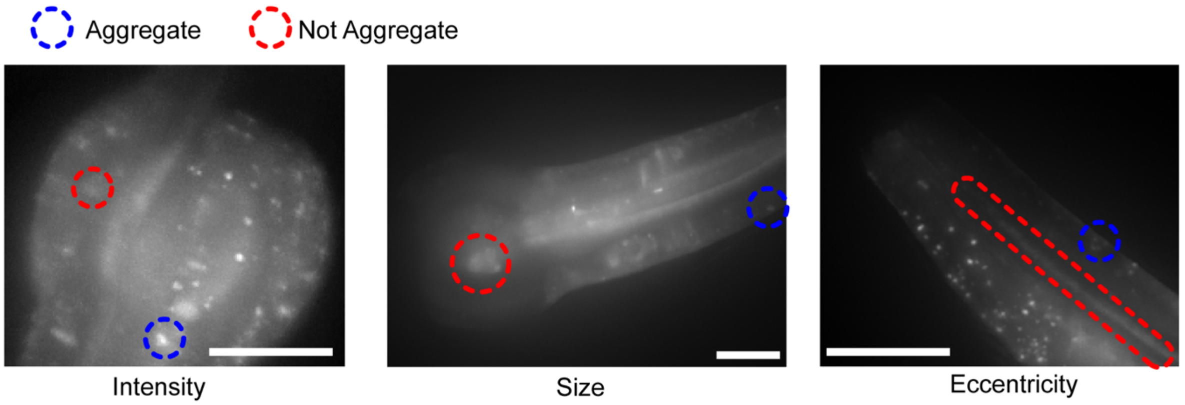


***Supplementary Figure 2*** *Examples of factors that differentiate aggregate objects from non-aggregate objects. Objects of certain size that are identified as aggregates exceed an intensity threshold relative to diffuse intensity. Objects of a certain intensity must be below a size threshold. Objects with high eccentricity are not identified as aggregates, as they are likely artefacts created by muscle fibers. Scale bar 10 µm.*

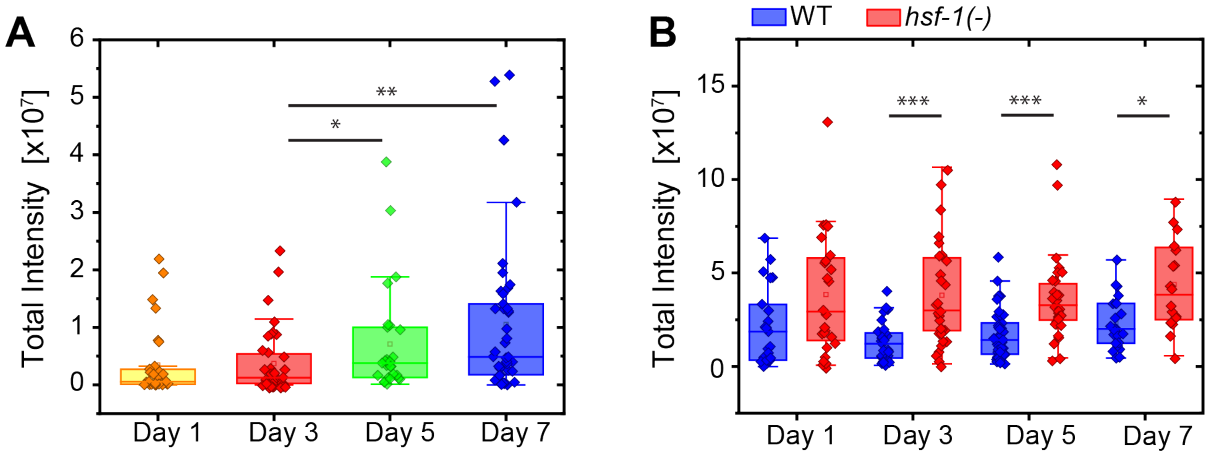


***Supplementary Figure 3*** *Increased levels of protein aggregation observed with age and in hsf-1 (-) mutants, quantified with the screening image processing algorithm.* ***(A)*** *Increase in total aggregate intensity observed from Day 1 to Day 7 of adulthood.* ***(B)*** *Increase in total protein aggregation intensity observed in hsf-1 (sy411) mutant relative to wild-type transgenic strain ,**p<0.01,**p<0.001, ***p<1E-4.*

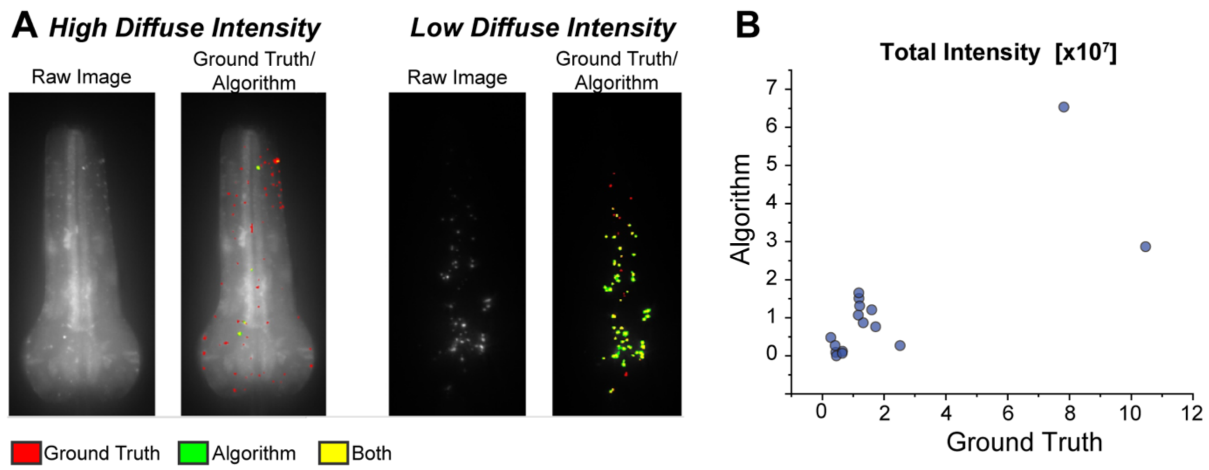


***Supplementary Figure 4 (A)*** *Sample images showing the low rate of aggregate identification in images with high diffuse intensity vs. low diffuse intensity. Ground Truth/Algorithm overlay ground truth binary (red) with algorithm binary (green). Overlap (yellow) is much higher in image with low diffuse intensity.* ***(B)*** *Predicted total intensity from algorithm plotted against actual total intensity from ground truth images.* $R^2$*=0.59.*

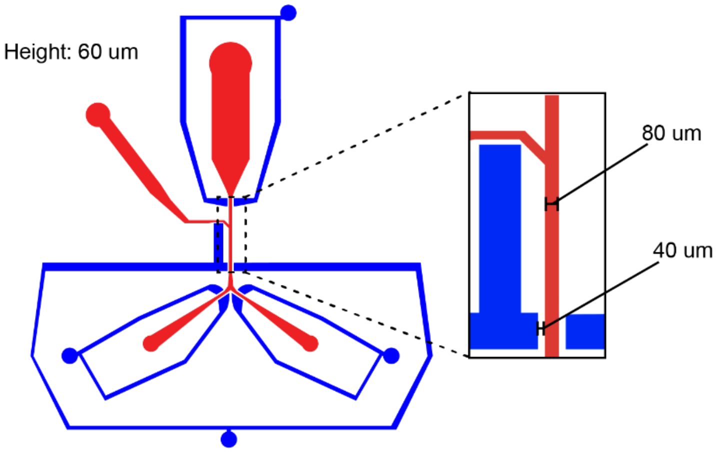


***Supplementary Figure 5*** *Dimensions of scaled-up microfluidic device for automated sorting. The feature height on the device is about 60 µm. The imaging channel width was widened to 80 µm, and the gap between the valve and channel was widened to about 40 µm.*

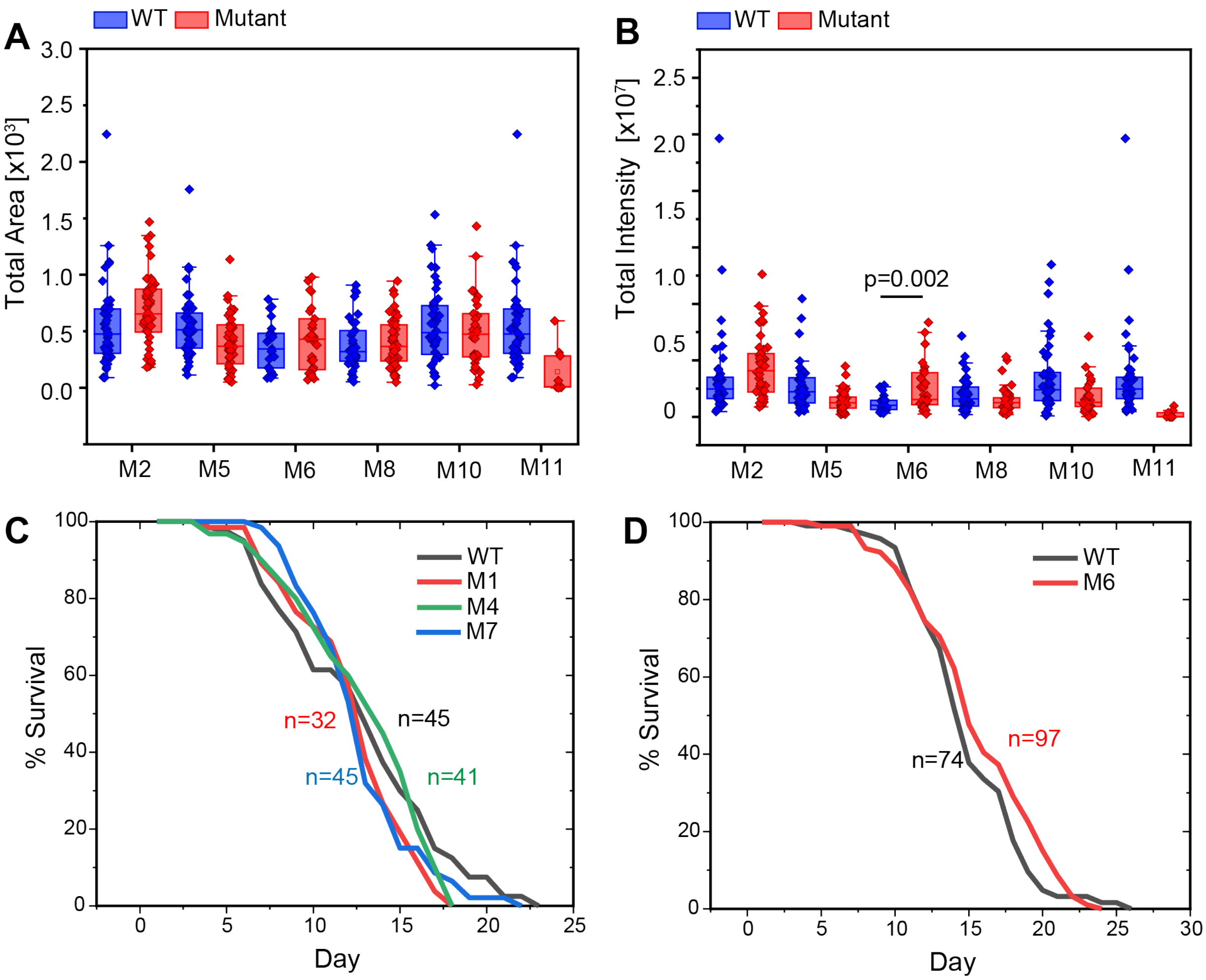


***Supplementary Figure 6 (A)*** *Total area and* ***(B)*** *total intensity measurements for mutants that do not show a significant increase in aggregation. Numbers under box plots indicate number of analyzed images* ***(C, D)*** *Lifespan curves for identified aggregation mutants that did not display a reduction in mean population lifespan. Mutant 6 was also tested for lifespan given its p-value close to significance (as per Bonferroni correction, $\alpha$=0.009)*

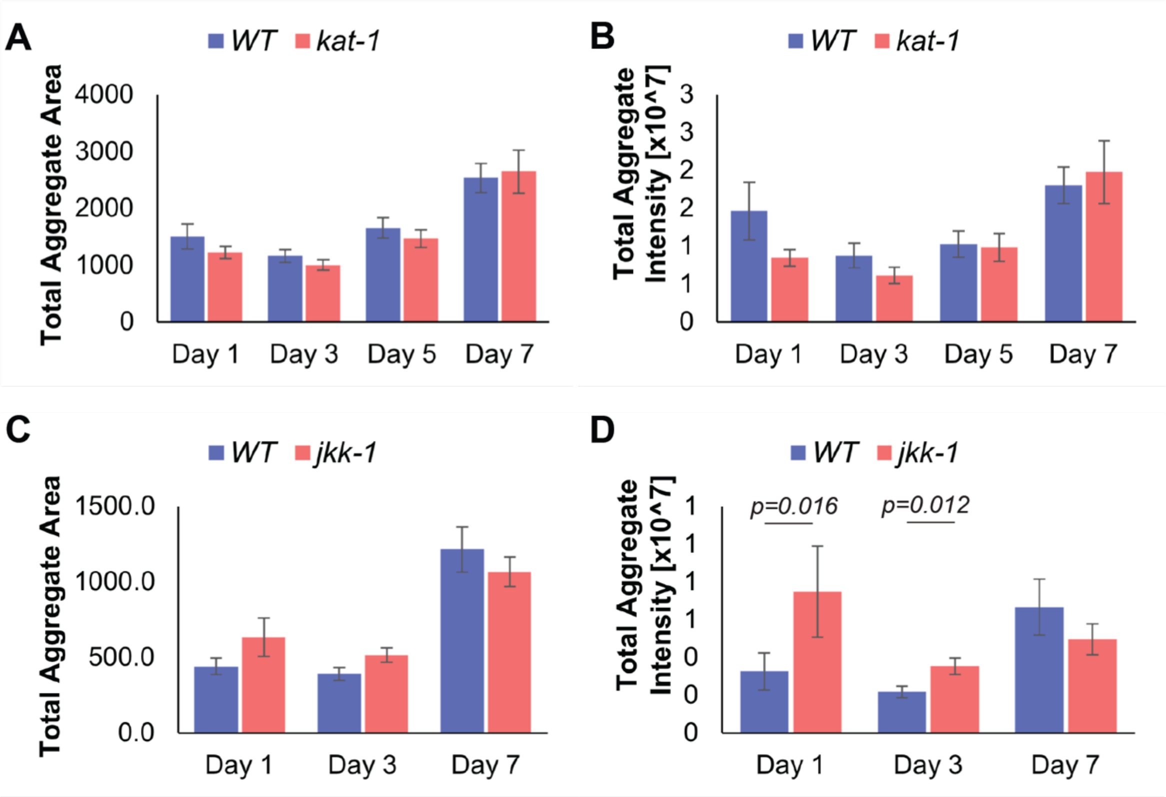


***Supplementary Figure 7*** *Total area and total intensity measurements for kat-1 and jkk-1 mutants, known to have a shortened lifespan. Error bars are S.E.M. p-values using Mann Whitney U Test*

***Supplementary Table 1***

| Feature Number | Description |
|---|---|
| **1** | Raw Pixel Intensity |
| **2** | Gradient Filter |
| **3** | Difference of Gaussian Filter 1 |
| **4** | Difference of Gaussian Filter 1 |
| **5** | Standard Deviation |
| **6** | Laplacian of Gaussian Filter |
| **7** | Disk Filter |
| **8** | Pixels Scaled to Maximum Image Intensity |
| **9** | Background Subtracted Feature 1 |
| **10** | Background Subtraction Feature 2 |
| **11** | Smoothed Standard Deviation Feature 1 |
| **12** | Smoothed Standard Deviation Feature 2 |
| **13** | Range Filter 1 |
| **14** | Range Filter 2 |

***Supplementary Table 2** Initial aggregation measurements acquired for putative total area and total intensity mutants, ranked by z-score.*

| Total Aggregate Area Z-score | | | | Total Aggregate Intensity Z-score | | |
|---|---|---|---|---|---|---|
| **1** | Mutant 1 | 5.85 | | **1** | Mutant 1 | 8.18 |
| **2** | Mutant 4 | 5.64 | | **2** | Mutant 6 | 4.78 |
| **3** | Mutant 3 | 5.33 | | **3** | Mutant 8 | 4.52 |
| **4** | Mutant 5 | 4.59 | | **4** | Mutant 11 | 4.23 |
| **5** | Mutant 6 | 4.49 | | **5** | Mutant 5 | 3.99 |
| **6** | Mutant 7 | 4.35 | | **6** | Mutant 3 | 3.99 |
| **7** | Mutant 8 | 4.13 | | **7** | Mutant 9 | 3.73 |
| **8** | Mutant 9 | 4.03 | | **8** | Mutant 2 | 3.2 |
| **9** | Mutant 10 | 3.81 | | **9** | Mutant 10 | 3.1 |
| **10** | Mutant 11 | 3.65 | | **10** | Mutant 7 | 2.91 |
| **11** | Mutant 2 | 3.52 | | **11** | Mutant 19 | 2.85 |
| **12** | Mutant 12 | 3.06 | | **12** | Mutant 4 | 2.75 |
| **13** | Mutant 13 | 2.91 | | **13** | Mutant 12 | 2.62 |
| **14** | Mutant 16 | 2.75 | | **14** | Mutant 14 | 2.45 |
| **15** | Mutant 14 | 2.55 | | **15** | Mutant 16 | 2.25 |
| **16** | Mutant 15 | 2.2 | | **16** | Mutant 18 | 2.15 |
| **17** | Mutant 17 | 2.03 | | **17** | Mutant 13 | 1.95 |
| **18** | Mutant 18 | 1.55 | | **18** | Mutant 15 | 1.54 |
| **19** | Mutant 19 | 1.46 | | **19** | Mutant 20 | 1.2 |
| **20** | Mutant 20 | 1.27 | | **20** | Mutant 17 | 0.76 |
| **21** | Mutant 27 | 1.27 | | **21** | Mutant 23 | 0.15 |
| **22** | Mutant 21 | 0.86 | | **22** | Mutant 27 | -0.67 |
| **23** | Mutant 22 | 0.55 | | **23** | Mutant 21 | -1 |
| **24** | Mutant 23 | 0.47 | | **24** | Mutant 24 | -1.11 |
| **25** | Mutant 24 | -0.02 | | **25** | Mutant 22 | -1.65 |
| **26** | Mutant 25 | -0.76 | | **26** | Mutant 25 | -1.87 |
| **27** | Mutant 26 | -1.76 | | **27** | Mutant 26 | -3.17 |
